# Inhibition of CDK7, CDK9 or CDK12 affects phosphorylation of human SPT5

**DOI:** 10.64898/2026.10.26.761550

**Authors:** Shona Murphy, Stephanie Barker, Girish R. Mali, Joey Riepsaame, Sophie Sheldon, Isabella Maudlin

## Abstract

SPT5, complexed with SPT4 in the DRB-sensitivity-inducing factor, DSIF, is a critical regulator of transcription elongation that acts negatively in the unphosphorylated form and positively when phosphorylated. *In vivo*, phosphorylation of multiple sites in the largely unstructured linker and C-terminal regions of the protein facilitates release from an early elongation checkpoint and processive transcription elongation. Transcription elongation factors, including the PAF complex (PAF1C), interact with phosphorylated SPT5 and phosphorylation of residues S666 and T806 are associated with the positive elongation functions of SPT5. Here, using *in vitro* phosphorylation of regions of the SPT5 C-terminus followed by GST-mediated pull-down we confirm that the RTF1 subunit of PAF1C interacts with residues 750-850 only when phosphorylated. Antibodies we raised to phospho-S666 or phospho-T806 are specific for these sites phosphorylated *in vitro* and *in vivo*. Western analysis of endogenous SPT5 in HEK293 cells using these antibodies before and after inhibition of analog-sensitive versions of CDK7, CDK9 and CDK12 indicate that all three kinases play roles in the phosphorylation of SPT5, although inhibition of CDK9 has the most marked effect. CDK9 inhibition also inhibits the association of PAF1C subunits with SPT5 *in vivo* and affects phosphorylation of chromatin-associated SPT5 as measured by chromatin immunoprecipitation (ChIP).

**Graphic abstract:** **CDK7, CDK9 and CDK12 regulate phosphorylation of SPT5**

CDK7, CDK9 and CDK12 regulate phosphorylation of the linker and CTR1 regions of SPT to facilitate recruitment of elongation factors. Inhibition of CDK7 mainly affects S666 phosphorylation, inhibition of CDK12 somewhat affects T806 phosphorylation, whereas inhibition of CDK9 greatly affects both marks, implicating all three kinases in the final pattern of SPT phosphorylation. The thickness of the arrows indicates the extent of the contribution to phosphorylation.

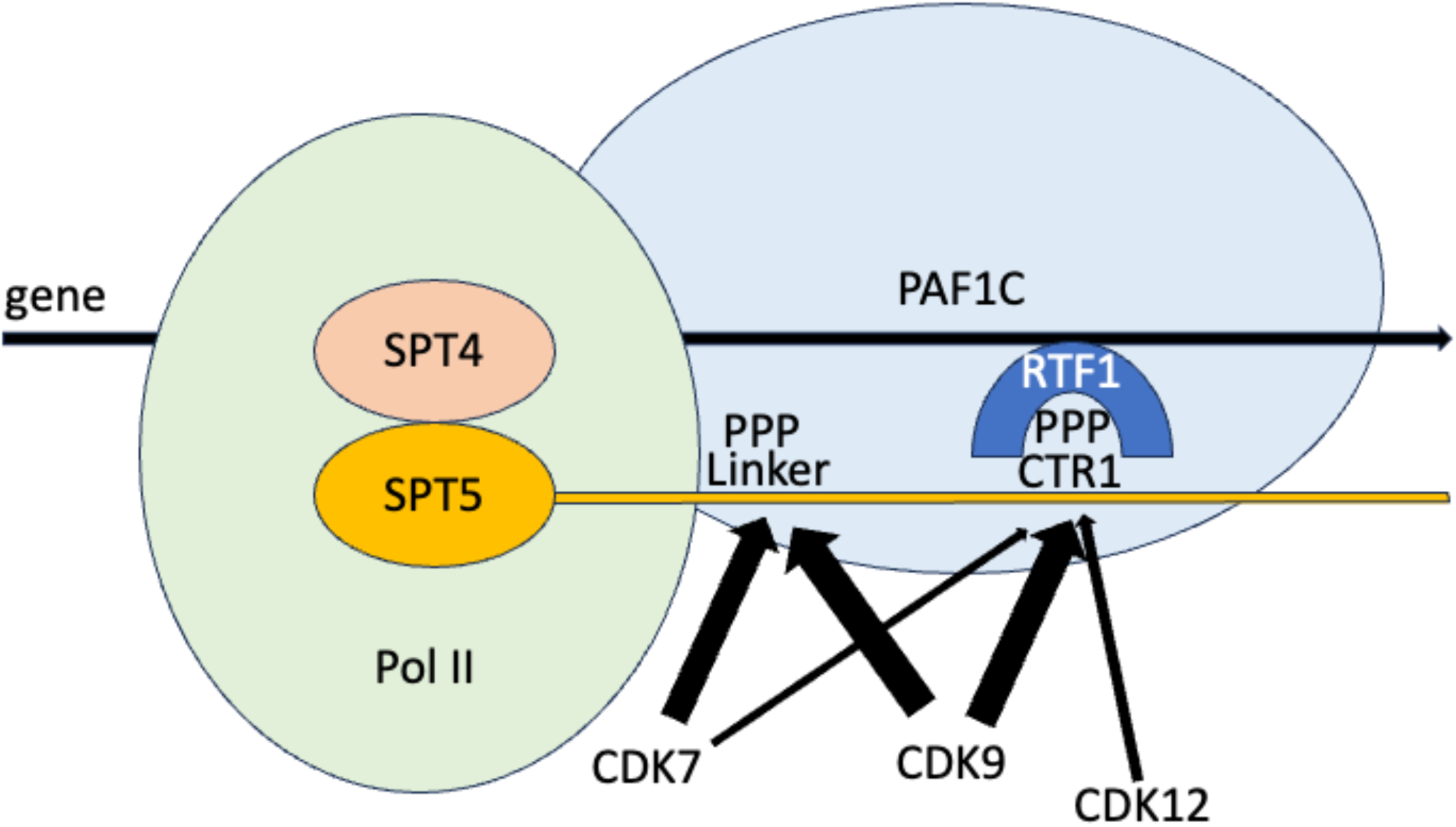

## Introduction

Elongation of transcription by RNA polymerase II (Pol II) is a highly-regulated process, involving a range of negatively- or positively-acting elongation factors[1–4]. Reversible phosphorylation of the carboxyl-terminal domain of the large subunit of Pol II (CTD) and several elongation factors, including SPT5, plays key roles in regulating transcription elongation[4–7].

SPT5 has homologues in all domains of life [4, 5, 7–9] and together with its partner SPT4 makes up the DRB-sensitivity-inducing factor (DSIF), which regulates transcription elongation[10]. In humans, SPT4 is a small (approximately 13 kDa), zinc finger-containing protein whereas SPT5 is a large (>120kDa) and complex protein comprising a structured amino-terminal acidic domain, a NusG N-terminal homology (NGN) domain, followed by seven Kyrpides-Ouzounis-Woese (KOW) domains and two largely unstructured C-terminal repeat regions; CTR1 and CTR2 (Figure 1)[4, 5, 7, 8, 11, 12]. The NGN domain of SPT5 interacts with SPT4 and Pol II[13], the KOW domains interact with Pol II, nucleic acid and NELF[14–17]. CTR1, CTR2 and the linker between KOW4 and KOW5 contain several potential phosphorylation sites for cyclin-dependent kinases (CDKs) (S/T-P) (Figure 1).

**Figure 1.**
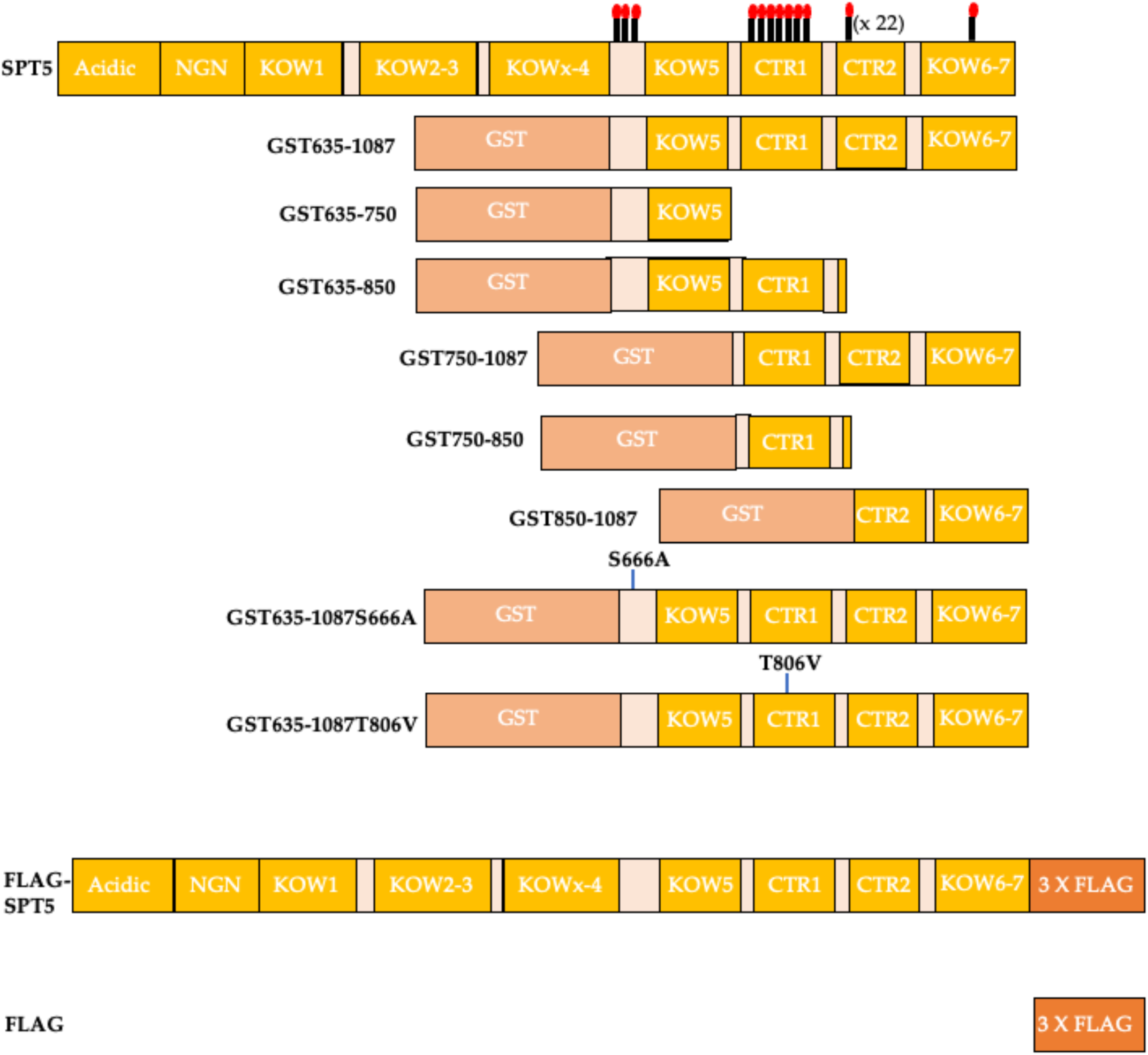
Schematic of the structure of human SPT5 and the constructs used in this study. The lollipops with a red head indicate potential S/T-P sites of phosphorylation by CDKs in the linker between the KOW4 and KOW5 domains, CTR1 (GS(Q/R)TPXY consensus TP sites), CTR2 (see[12]) and further towards the C-terminus (see https://www.phosphosite.org for reported *in vivo* phosphorylation). The GST-tagged proteins were produced in *E.coli* and the FLAG-tagged proteins were produced in transfected CDK9as HEK293 cells.

DSIF joins the Pol II elongation complex soon after initiation and together with the negative elongation factor, NELF, helps to pause Pol II close to the promoter[18–20]. Phosphorylation of NELF by the cyclin-dependent kinase 9 (CDK9) subunit of positive transcription elongation factor b (P-TEFb) destabilizes its interaction with the Pol II complex and phosphorylation of SPT5 facilitates the binding of the polymerase-associated factor 1 (PAF1) elongation complex (PAF1C) to Pol II to displace NELF and allow productive elongation[18–20]. SPT5 is therefore unusual in that it functions as a negative elongation factor when unphosphorylated and a positive elongation factor when residues in the linker between KOW4 and 5 and the C-terminal unstructured region are phosphorylated. SPT5 also plays roles in Pol II stability on genes and enhancers[21–23], RNA processing [5, 24] and termination of transcription[25–29].

Several cyclin-dependent kinases regulate the transcription cycle[6, 30–32]. CDK7, a subunit of the general initiation factor TFIIH, phosphorylates the Pol II CTD and regulates promoter escape[30, 33–35]; CDK9 is a master regulator, controlling escape from the promoter-proximal pause, termination of transcription, RNA processing and histone modification[2, 6, 12, 19, 36–38]; CDK12 phosphorylates the CTD of Pol II and activates elongation of transcription, at least in part through phosphorylation of the Leo1 subunit of PAF1C and recruitment of PAF1C subunits and SPT6, another positive elongation factor, to the Pol II elongation complex[39, 40]. Accordingly, CDK12 plays a critical role in expression of long genes[41, 42]. CDK12 shows redundancy with the highly-related CDK13 and inhibition of both causes major elongation defects[43].

SPT5 is thought to be largely phosphorylated by CDK9 [12, 44] and CDK9 inhibitors affect SPT5 phosphorylation in cells[12, 29, 38, 45]. For example, phosphorylation of serine (S)666 and S671 in the linker between KOW4 and KOW5 is decreased by the CDK9 inhibitor, DRB[38]. In addition, S666 and threonine (T)806, in CTR1, are targets of *in vitro* phosphorylation by CDK9[46]. CDK7 can also phosphorylate S666[34]. Using phospho-specific antibodies it has been shown that phospho-(p)S666 largely follows the pattern of SPT5 across genes, whereas pT806 decreases after the poly(A) site[29]. Phosphorylation of mammalian SPT5 is reversed by PP1, PP2A as part of INTAC and PP4[27, 29, 47, 48].

In line with the role of SPT5 phosphorylation in control of elongation, mutation of S666, S671 and S686 to alanine impairs release of the promoter-proximal pause [12, 23, 49] and phosphomimetic substitutions in the CTR1 domain are associated with decreased pausing[49]. CTR2 phosphorylation instead seems to have a braking effect on elongation[12]. *In vitro* interaction analysis using recombinant protein and an SPT5 CTR1 phospho-peptide comprising one of the seven consensus T-P repeats (see Figure 1) indicates that phosphorylated threonine residues in CTR1 interact with the Plus3 domain of the RTF1 subunit of PAF1C[50]. In addition, chromatin association of the RTF1 subunit of PAF1C is decreased when CTR1 cannot be phosphorylated[12].

Here, we show that, *in vitro*, the phosphorylated CTR1 region of human SPT5 is sufficient for interaction with full-length RTF1 in nuclear extract, whereas the linker region or CTR2 are not. Antibodies raised to pS666 specifically recognize the *in vitro* phosphorylated linker region and antibodies raised to pT806 specifically recognize the *in vitro* phosphorylated CTR1. Western analyses using these antibodies indicates that specific inhibition of CDK7, CDK9 or CDK12 in cells where the endogenous kinases are analog-sensitive (as) affects SPT5 phosphorylation. However, inhibition of CDK9as causes the greatest loss of phosphorylation of both pS666 and pT806. Flag-mediated pull down of SPT5 from CDK9as cells confirms that association of PAF1C subunits, PAF1, RTF1 and CDC73 with SPT5 is decreased after inhibition of CDK9as. In addition, chromatin immunoprecipitation (ChIP) analysis indicates that pS666 and pT806 both decrease markedly at the beginning of the model *KPNB1* gene when CDK9as is inhibited.

These results underscore the critical role of cyclin-dependent kinases in SPT5 phosphorylation and confirm the key role of CDK9 in recruitment of the PAF1C complex to the Pol II elongation complex through SPT5 phosphorylation.

## Materials and Methods

### Production of GST-tagged SPT5 constructs

The regions of SPT5 outlined in Figure 1 were cloned into pGEX-4T-1 (Merck) using the BamH1 and Not1 restriction sites. The plasmids were transformed into Rosetta DE3 bacteria and the GST-tagged SPT5 proteins were produced and purified as detailed [51] using glutathione magnetic agarose beads (Pierce) (Thermo Fisher 78601). Following elution, the glutathione was removed using Zeba spin desalting columns (Thermo Fisher 89882) before the concentration was measured by SDS-PAGE against BSA standards.

### In vitro phosphorylation of the GST-SPT5 proteins

Equivalent molar amounts of GST, GST-635-1087, GST-635-750, GST-750-850 or GST-850-1087 were incubated with 10μg HeLa nuclear extract in 100μl final volume o/n at 30°C in 20mM Tris pH8, 8mM MgCl2, 50mM KCl, 5mM β-glycerol phosphate, 20mM creatine phosphate, 1mM ATP, 1x PhosSTOP, (Merck 4906845001), cOmplete™, EDTA-free Protease Inhibitor Cocktail, Sigma-Aldrich. Control reactions contained BC100 [52] instead of nuclear extract.

### Pull down of GST-SPT5 proteins from HeLa nuclear extract

For each pull down, 20 μl of each phosphorylation (or control) reaction and 80 μl 20mM Tris pH 8.0, 0.2mM EDTA, 50mM KCl was added to dry 5μl glutathione magnetic agarose beads (washed in 20mM Tris pH 8.0, 0.2mM EDTA, 50mM KCl) and incubated on a wheel at 4°C for 90 minutes to capture the GST-tagged proteins. The beads were washed 2x with 20mM Tris pH 8.0, 0.2mM EDTA, 300mM KCl and 1x with 20mM Tris pH 8.0, 0.2mM EDTA, 50mM KCl. The dry beads were resuspended in 100μl buffer with a final concentration of 20mM Tris pH8, 2.5mM MgCl2, 50mM KCl, 0.5% Igepal (Merck I8896), 4% glycerol, 1x PhoSTOP, 1x Pierce protease inhibitors and 200μg HeLa nuclear extract and incubated o/n at 4°C on a wheel. The supernatant was collected and the beads were washed 2x with 20mM Tris pH 8.0, 0.2mM EDTA, 300mM KCl, 1x PhoSTOP, protease inhibitors, 1x with 20mM Tris pH 8.0, 0.2mM EDTA, 50mM KCl, 1x PhoSTOP, protease inhibitors. Bound proteins were eluted by heating the beads with LDS sample buffer with 100mM DTT.

### Generation of phosphoantibodies

Rabbit phosphoantibodies against SPT5 pS666 or pT806 were generated by Eurogentec based on the peptide sequences: pS666 (663-674) H-C-APM S(PO3H2) PRISSPMH-NH2; pT806 (799-813) H-TPHYGSQ T(PO3H2) PLHDGSR-NH2.

Phosphoantibody specificity was ensured by selecting against recognition of the unphosphorylated peptide and by lack of reaction to the recombinant GST-SPT5 proteins with the relevant amino acid replaced by a non-phospho-acceptor (Figure 2c).

**Figure 2.**
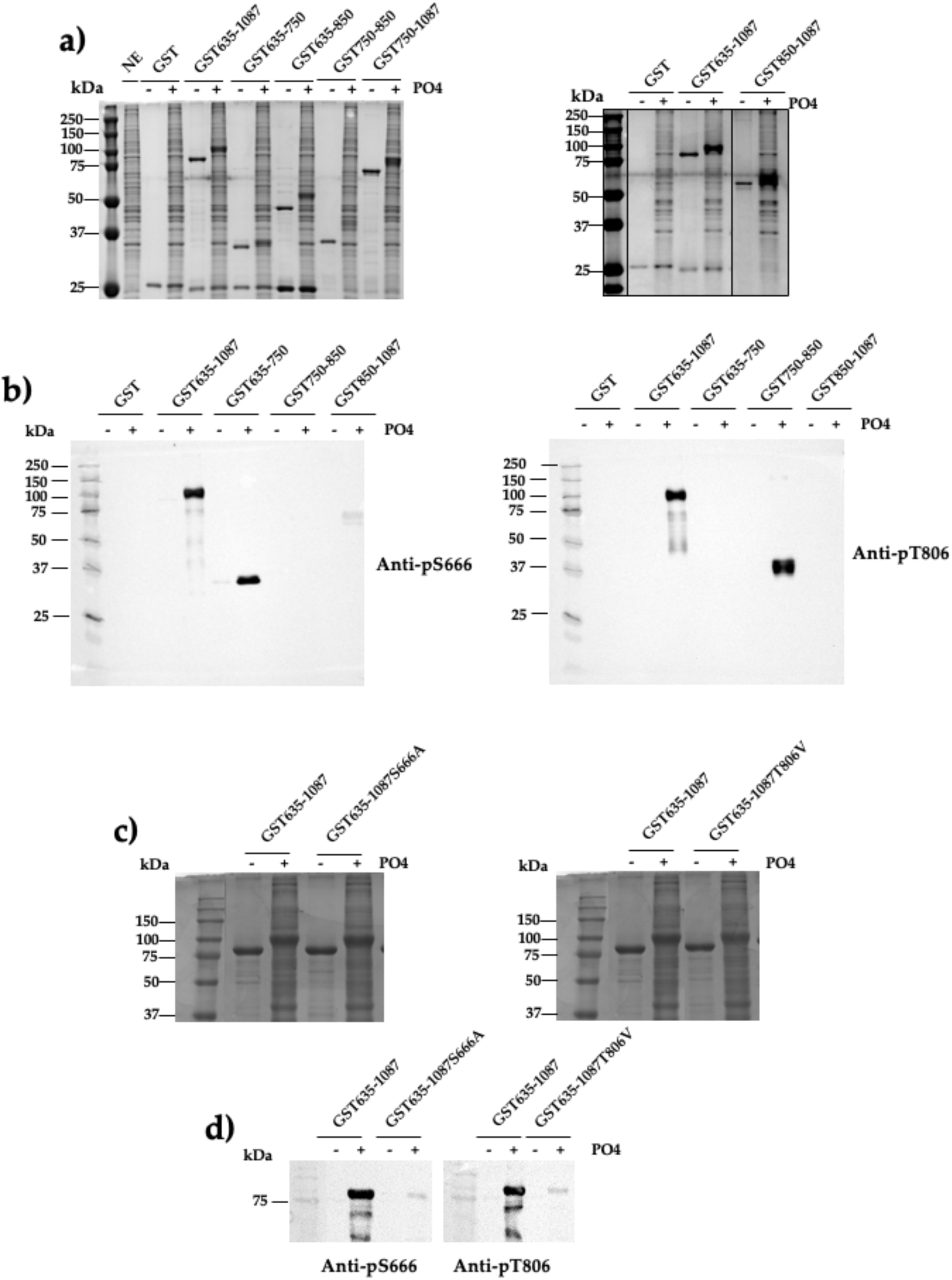
The SPT5 C-terminus can be phosphorylated *in vitro*. **a)** Coomassie-stained 10% SDS-PAGE gels of GST-SPT5 proteins produced in *E.coli* using constructs depicted in Figure 1. The protein and phosphorylation status are indicated at the top and the sizes of the molecular weight markers are indicated at the left. **b)** Western blot analysis of GST-SPT5 proteins with anti-pS666 and anti-pT806 antibodies as indicated on the right. The protein and phosphorylation status are indicated at the top and the sizes of the molecular weight markers are indicated at the left. **c)** Coomassie-stained 10% SDS-PAGE gels of GST-SPT5 proteins produced in *E. coli* using constructs depicted in Figure 1. The protein and phosphorylation status are indicated at the top and the sizes of the molecular weight markers are indicated at the left. **d)** Western blot analysis of GST-SPT5 proteins with anti-pS666 and anti-pT806 antibodies as indicated on the right. The protein and phosphorylation status are indicated at the top and the sizes of the molecular weight markers are indicated at the left. NE= nuclear extract.

### HEK293 CDKAS Cell Culture

All human embryonic kidney (HEK) 293 CDKas cells were grown in Dulbecco’s Modified Eagle’s Medium (DMEM) with 10% fetal bovine serum, 2 mM L-glutamine, 50 U/mL penicillin and 50 µg/mL streptomycin and incubated at 37°C in 5% CO2. The medium was changed 16-18 hours before adding inhibitors. Cells were treated with 10 µM 1-NM-PP1 (CDK7as and CDK12as or 15 µM 1-NA-PP1 (CDK9as) (Cayman Chemical Company) or 100 µM DRB (Abcam ab120939) for 30 minutes, as indicated.

### Western blotting

Western blot analysis was performed as previously described [53] using recombinant protein, nuclear lysate or pulldown eluate resuspended in 1x Pierce LDS sample buffer (Thermo Fisher 84788) with 100mM DTT and incubated for 10 minutes before loading on 6-10% PAGE gels with APEX bio prestained protein marker (F4005). Gels were run and wet transferred to nitrocellulose (Thermo Fisher 88018) at 4°C in 10mM Tris base, 100mM Glycine, 10% ethanol o/n. Blots were imaged on an iBright FL1500 and quantitation was carried out using iBright software. The antibodies used are listed in Table 1. Secondary antibodies were purchased from Merck Millipore (Goat Anti-Rabbit IgG Antibody, HRP-conjugate, 12–348, and Goat Anti-Mouse IgG Antibody, HRP conjugate, 12–349), the chemiluminescent substrate (SuperSignal West Pico PLUS) from Thermo Fisher, and the membranes visualized on an iBright FL1000 Imaging System (Thermo Fisher). Quantification of the western blots was performed with iBright analysis software.

**Table 1.** List of antibodies used for Western blot or ChIP.

| Antibodies | Reference |
| --- | --- |
| Anti-Flag (M2) | Merck F3165 |
| Pol II N-terminus | Cell Signaling D8I4Y |
| pSer2 CTD | Abcam ab5095 |
| pSer5 CTD | Abcam ab 5131 |
| SPT5 | Santa Cruz H300 (western) or H300X (ChIP) |
| RTF1 | Abcam ab184570 |
| PAF1 | GTX107346 |
| Cdc73 | Thermo Fisher A300-170A |

It should be noted that RTF1 can break down if heated at 95oC in SDS sample buffer in the presence of Dynabeads.

### Chromatin immunoprecipitation (ChIP)

ChIP and qPCR were performed essentially as described [54] using approximately 1 x 10⁷ HEK293 CDK9as cells cross-linked with 1% formaldehyde at room temperature for 10 min after treatment of the cells with DMSO or inhibitor. For real-time qPCR, ChIP samples were analysed using the QuantiTect SYBR Green PCR kit (Qiagen) and Rotor-Gene RG-3000 (Corbett Research). Data presented represent the mean percentage (%) of Input ± SEM (standard error of the mean) of three independent experiments. The primers used for qPCR are described in [39]. The antibodies used are listed in Table 1.

### Analog-sensitive CDK12 (CDK12as) cell line creation

Cas12a ULTRA protein, Alt-R Cas12a electroporation enhancer, ssODN HDR templates, tracrRNA and synthetic crRNAs were purchased from Integrated DNA Technologies (IDT). The Neon Transfection System (cat. #MPK5000) and consumables/buffers (cat. #MPK1096) were purchased from Thermo Fisher Scientific. For Cas12a RNP formation, 2 ug Cas12a ULTRA protein (IDT) was mixed with 15 pmol of each crRNA in a total volume of 2 ul IDTE Buffer (IDT, cat. # 11-05-01-04), resuspended 5 times and incubated at 37°C for 5 minutes. Resulting RNP mixes were added directly to cells prior to nucleofection.

HEK293 were washed twice with PBS and resuspended in 8 uL “buffer R”. Subsequently, cells were mixed with 2 uL RNP solution from the previous step, 2 uL of a 2.16 uM Alt-R Cas12a electroporation enhancer (IDT, cat. # 1076300, diluted in IDTE buffer) and 125 pmol ssODN HDR template. This mixture was resuspended five times in a 10 uL Neon nucleofector tip and nucleofected with the Neon Transfection System using the following settings: 1700 V, 20 ms, 1 pulse (program “J”) or 1325 V, 10 ms, 3 pulses. Upon nucleofection, cells were transferred directly into a single well of a 24w plate containing 500 uL pre-warmed complete medium and allowed to recover overnight in a humidified incubator at 37°C under 5% CO2 atmosphere. Medium was replaced with fresh complete medium the next morning and cells were allowed to expand to 70-80%. Guides: crRNA1 5’-AGTATATGGACCATGACTT-3’; crRNA2 5’-CCTTGTATTTGAGTATATA-3’. ssODN (HDR template) (TTT-to-GGC) 5’-T*T*AATTTTTTTTCGTCTTTATGTAGGTGCCTTTTACCTTGTAGGCGAGTATATGGAC-CATGACTTAATGGGACTGCTAGAATCTGGTT*T*G3’, where the asterisks denote phosphorothioate modification[55]. For screening, **c**ells were incubated in 180 µl ChIP lysis buffer for 10 min and sonicated for 3 min (30 s on-30 s off) using a Q800R2 sonicator (QSONICA). 20 µl ammonium acetate was added (final conc. 400 mM) and samples were mixed with 200 µl phenol chloroform by vortexing. The samples were centrifuged (13 000 g, 5 min, 25°C) and the upper phase was transferred to a new tube. Genomic DNA was precipitated in 70% ethanol, pelleted by centrifugation (13 000 g, 5 min, 25°C) and dissolved in nuclease free water. A CDK12 fragment was PCR amplified using the following primers: Forward 5′ CCAGCCATATGGGTGGCAGAGTGAGAGTCT 3′; Reverse 5′ CCAGCGCGGCCGCG-

CAGAAAAGGCCAATCA 3′. DNA was purified using a QIAquick PCR purification kit (Qiagen) and sequenced by the SourceBioscience Sanger Sequencing Service, Oxford (Supplementary Figure). RNA was extracted from HEK293 and CDK9as cells using a *Quick*-RNA Miniprep kit (Zymo Research) according to the manufacturer’s instructions. Reverse-transcription (RT) was performed with 500 ng of RNA using random hexamers with the SuperScript III kit (Invitrogen) according to the manufacturer’s instructions. Sequencing was performed on a cDNA PCR fragment generated with Forward primer: CGCTGTGTGGACAAGTTTGA and Reverse primer: GCTGGTGTGTAACGTTCCTC) (Supplementary Figure). Sequencing primer used: GCTTCCCAATCACAGCCATT.

### Cell proliferation analysis

Cells were seeded at 500/well in 95 µl medium in 96-well microplates (Greiner, 655090) and cell number was measured at the time points noted on Supplementary Figure b and c by adding alamarBlue HS (ThermoFisher, A50101) (1/20) and reading in a fluorimeter after 1 hour incubation, according to the manufacturer’s instructions. 1-NM-PP1 (NM) or DMSO was added to the cells as noted on Supplementary Figure c by an arrow.

### Co-immunoprecipitation of FLAG-tagged SPT5

Co-IP was performed essentially as detailed in[38]. For each sample, 10 µl of Dynabeads protein G (Thermo Fisher 10765583) were pre-blocked o/n at 4°C on a wheel at 16 rpm in 0.5 ml of TBS with 0.5% BSA and washed three times in IP buffer (25 mM Tris–HCl pH 8.0, 150 mM NaCl, 0.5% NP-40, 2.5 mM MgCl2), before being incubated o/n at 4°C on a wheel at 16 rpm in 300 µl of IP buffer with 1 µg of anti-FLAG antibody, protease inhibitors and PhosSTOP. The beads conjugated with antibodies were washed three times with IP buffer supplemented with PhosSTOP and protease inhibitors cocktail and incubated with approximately 0.2 mg of protein at 4°C on a wheel at 16 rpm o/n. 1 x 15cm dish of 60– 70% cells was transfected with SPT5-FLAG (full length SPT5 in pCDNA3 with 3XFLAG at C-terminus) or FLAG (3XFLAG in pCDNA3) as control using jetPrime (Polyplus), changing the medium after 5 hours. 16-18 hrs later, the cells were treated with DMSO or inhibitor for 30 minutes and nuclei were prepared as detailed[53]. Nuclei were lysed by adding 400 μl lysis buffer (50mM Tris pH 8.0, 150mM NaCl, 1% Igepal, 2.5mM MgCl2, 1x PhosSTOP, 1x protease inhibitors), approximately 25 units of Benzonase (Merck 1.01654) was added and the samples were incubated at 4°C on a wheel for 1 hour. Samples were spun at 4K RPM for 1 minute at 4°C and the supernatant collected. An equal volume of dilution buffer (150mM NaCl, 2.5mM MgCl2, 1x PhosSTOP, 1x protease inhibitors) was added and this was the input sample. Protein concentration was determined by Bradford assay. After incubation of the nuclear lysate with the FLAG antibody-charged Dynabeads, the beads were washed three times with IP buffer with PhosSTOP and protease inhibitors and three times with IP buffer with PhosSTOP and protease inhibitors without Igepal. Proteins were eluted in 1× LDS plus 100 mM DTT for 10 min at 70°C. Proteins were separated on 6% SDS PAGE gels and transferred to nitrocellulose o/n at 4oC in a wet transfer apparatus as detailed above.

### AlphaFold structure prediction and modelling

AlphaFold 3 (AF3) [56](AF server: https://alphafoldserver.com/) was used for predicting and modeling potential interactions between SPT5 and RTF1 in the absence and presence of phosphorylation. Model statistics (predicted alignment error plots) are as generated by the algorithm with annotations to highlight relevant details of protein boundaries and predicted interaction interfaces. Final modeling and visualizations were performed using UCSF ChimeraX[57].

## Results

### The SPT5 C-terminus can be phosphorylated in vitro

SPT5 is phosphorylated on residues S666, S671 and S686 in the linker between the KOW4 and KOW5 domains, in CTR1 and CTR2 and downstream of CTR2 *in vivo* [27, 29, 38] (https://www.phosphosite.org) (Figure 1). Mutational studies and ChIP with antibodies to pS666 and pT806 implicate these phosphorylated residues in regulation of elongation[12, 23, 27, 29, 49]. We have cloned residues 635-1087, 635-750, 635-850, 750-850, 750-1087 and 850-1087 of SPT5 into pGEX-4T to enable production, in bacteria, of C-terminal GST-tagged proteins including these residues (GST635-1084, GST635-750, GST635-850, GST750-850, GST750-1087 and GST850-1087) and residues 635-1087 with either S666 mutated to alanine or T806 mutated to valine (GST635-1087S666A and GST635-1087&806V) (Figures 1, 2a, 2c).

All GST-tagged proteins have retarded mobility after incubation with HeLa nuclear extract in conditions for phosphorylation (Figure 2a, 2c), suggesting that phosphorylation has occurred. Importantly, rabbit antibodies raised to phosphor-S666 or phosphor-T806 efficiently recognize only the appropriate phosphorylated proteins (Figure 2b). In addition, the antibody to pS666 does not efficiently recognize phosphorylated GST-tagged 635-750 when S666 is mutated to alanine and the antibody to pT806 does not efficiently recognize phosphorylated GST-tagged 750-850 when T806 is mutated to valine (Figure 2d), demonstrating their specificity.

These results indicate that some of the phospho-acceptor residues between 635 and 1087 of SPT5, including S666 and T806 can be phosphorylated *in vitro* and that antibodies raised to SPT5 peptides containing pS666 or pT806 are specific for the appropriate phospho-epitope.

### RTF1 binds to region 750-850 of SPT5 only when it is phosphorylated

It has already been shown that the RTF1 subunit of the PAF complex binds to a peptide comprising residues 779-790 of SPT5 when this is phosphorylated on residue T784[50]. We have therefore tested whether the GST635-1087, GST635-750, GST635-850; GST750-850, GST750-1087 and GST850-1087 proteins can bind to RTF1 before or after phosphorylation by performing GST-mediated pull down of these proteins incubated with HeLa nuclear extract, followed by western analysis (Figure 3a). RTF1 binds to GST635-1087, GST635-850, GST750-850 and GST750-1087 only when these proteins are phosphorylated, but not to GST635-750 or GST850-1087, whether these proteins are phosphorylated or not (Figure 3a, b, c), indicating that *in vitro* phosphorylation of SPT5 residues between 750 and 850 can facilitate the interaction between the C-terminus of SPT5 and RTF1.

**Figure 3.**
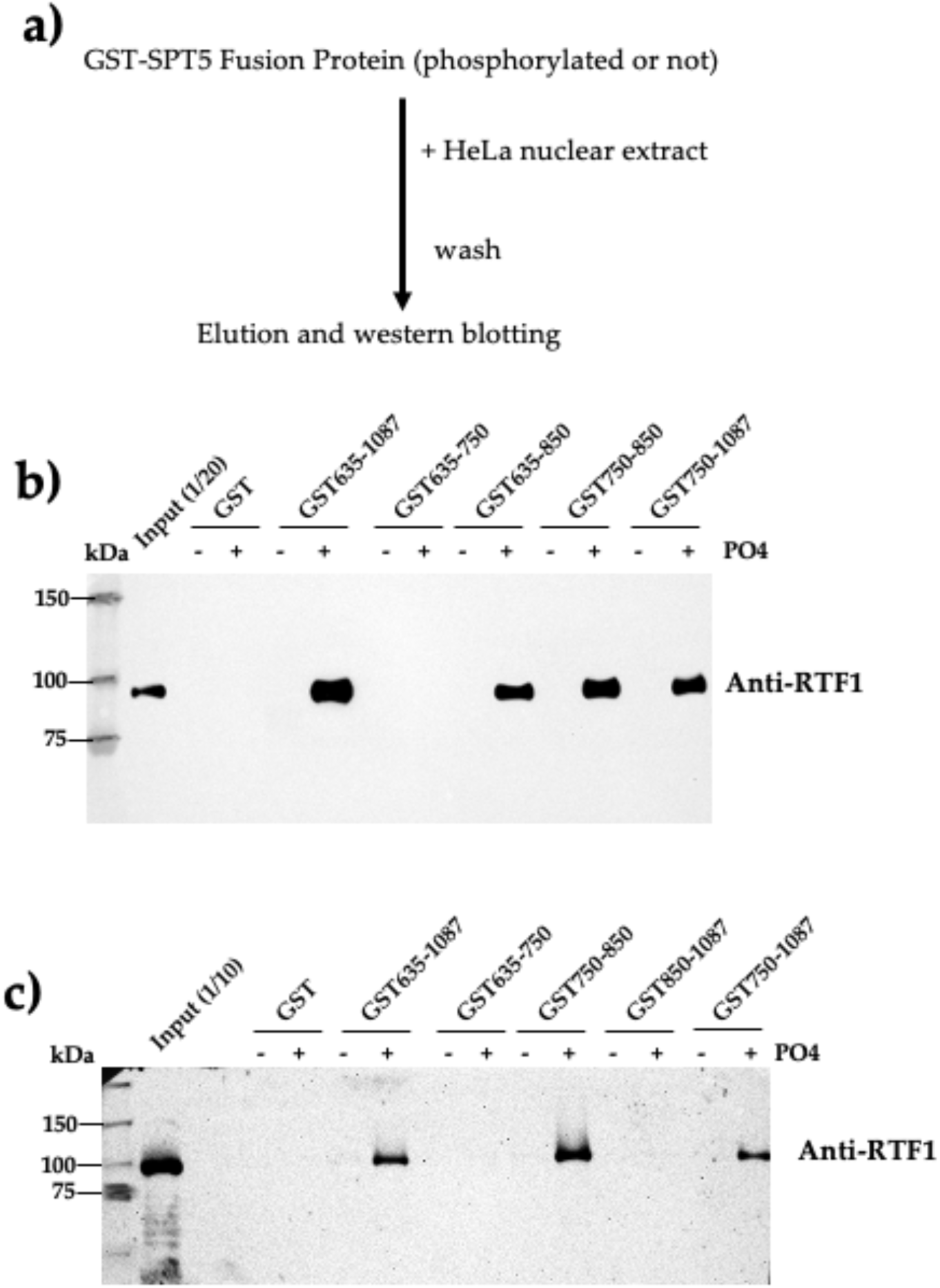
RTF1 binds to region 750-850 of SPT5 only when it is phosphorylated. **a)** Schematic of the pull-down procedure. **b), c)** Western blot analysis of the eluates of pull down of GST-SPT5 proteins with an anti-RTF1 antibody as indicated on the right. The protein used and phosphorylation status are indicated at the top and the sizes of the molecular weight markers are indicated at the left.

This recapitulates the interaction between a phosphorylated SPT5 peptide comprising residues 779-790 and RTF1 [50] and further indicates that other regions upstream or downstream of 750-850 in GST-635-1087 are not sufficient to bind to RTF1, even when phosphorylated. Western analysis was also carried out for other subunits of the PAF1C. However, the antibodies used recognize the GST-tagged proteins alone, whether incubated with extract or not (data not shown), making it difficult to determine if any other PAF1C subunits bind to the unphosphorylated or phosphorylated proteins.

### Alphafold models the interaction between RTF1 and SPT5

Alphafold 3 (https://alphafoldserver.com/) was used to model the interaction between SPT5 and RTF1 when SPT5 was either unphosphorylated or phosphorylated on T806 or on all seven GS(Q/R)TPXY consensus TP residues between 750 and 850 (CTR1) (Figure 4) (see Figure 1). No interaction is predicted when SPT5 is unphosphorylated (a), whereas some interaction is predicted when T806 of SPT5 alone is phosphorylated (b). In addition, Alphafold predicts a broader interaction interface between SPT5 and RTF1 when the seven threonine residues in CTR1 region are phosphorylated (c and d). This predicted model of the interaction between SPT5 phosphorylated in CTR1 and RTF1 is consistent with the results in Figure 3, those of Sun and Fisher (2025) [12]and those of Wier et al (2013), indicating that the Plus3 domain of RTF1 interacts with an SPT5 CTR1 phospho-peptide[50].

**Figure 4.**
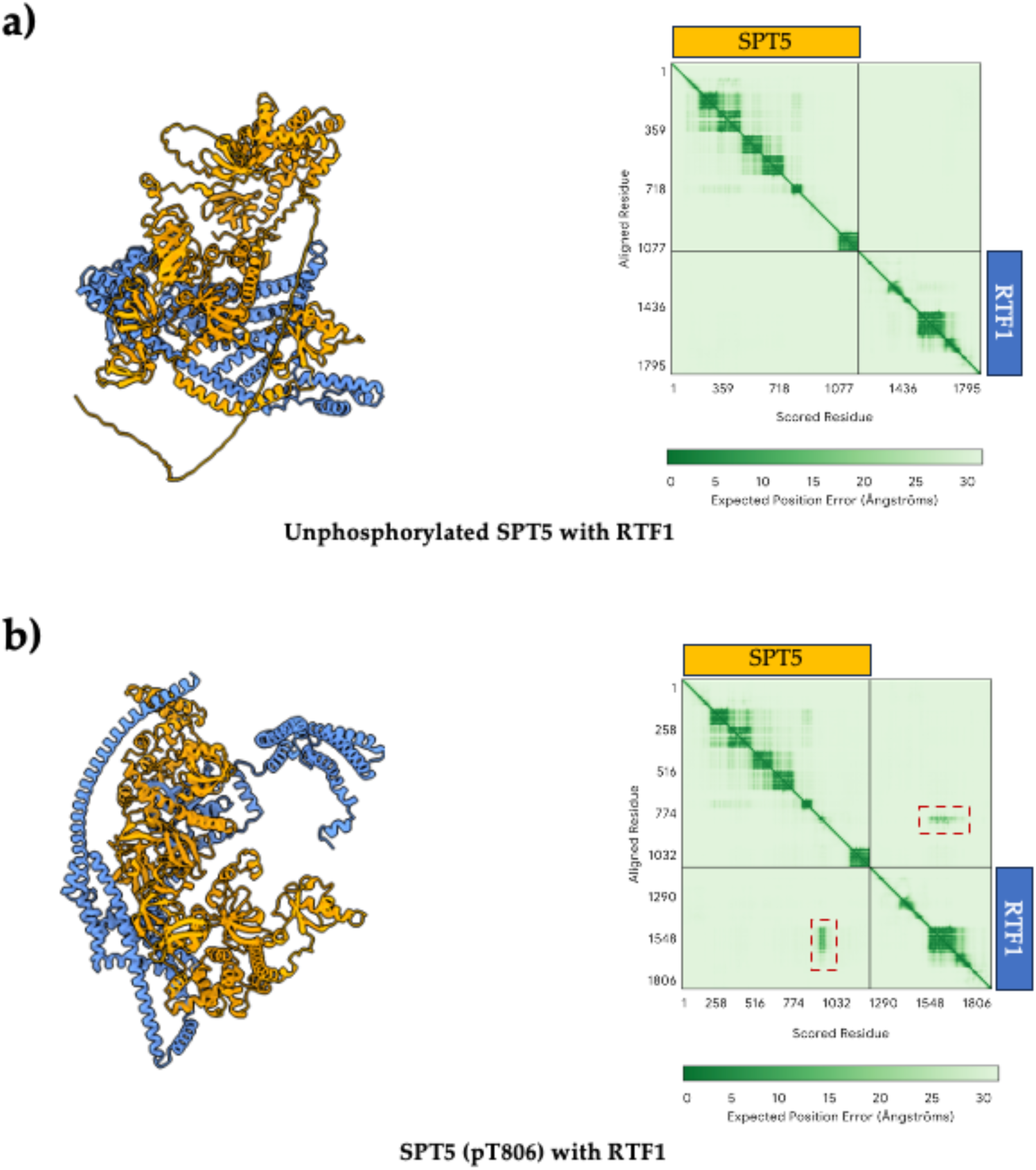
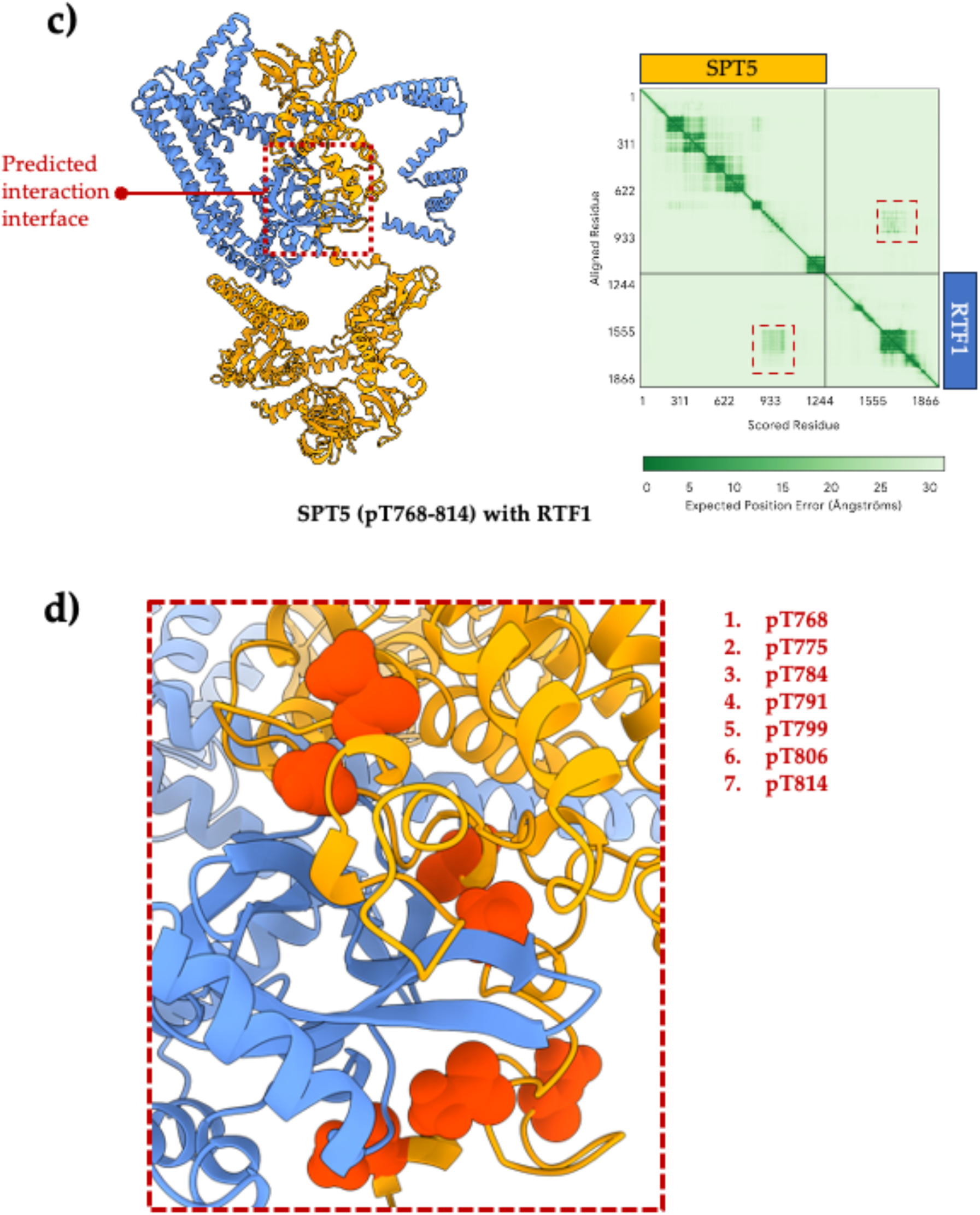
Alphafold models the interaction between RTF1 and SPT5. **a)-c)** AlphaFold predictions of complexes between full-length SPT5 [in orange; unphosphorylated in (a), phosphorylated at threonine 806 in (b) and phosphorylated at the seven threonines between 750 and 850 reported as phosphorylated *in vivo* (https://www.phosphosite.org) and conforming to the GS(Q/R)TPXY consensus (c) and full-length RTF1 (in blue) are shown with the associated Predicted Alignment Error (PAE) plots at the right. Red dashed boxes in the PAE in (b) and (c) highlight predicted interaction interfaces between SPT5 and RTF1. ipTM score in (a) = 0.15, in (b) = 0.25 and in (c) = 0.23. Note the lack of interaction between unphosphorylated SPT5 and RTF1 in (a). **d)** Predicted interaction interface between SPT5 and RTF1 highlighted in (c) is shown in closer detail. The seven phospho-threonines between 750 and 850 are depicted as red spheres on the SPT5 structure and the positions listed next to the panel.

### Inhibition of CDK7, CDK9 or CDK12 affects phosphorylation of SPT5 at S666 and T806

To assess the role of CDK7, CDK9 and CDK12 in phosphorylation of SPT5 *in vivo*, nuclei were prepared from CDK7as, CDK9as or CDK12as cells before or after treatment of cells with the appropriate analog (1-NM-PP1 for CDK7as and CDK12as cells or 1-NA-PP1 for CDK9as cells as detailed in Materials and Methods). Production of CDK7as and CDK9as cells has been described previously[35, 38]. The CDK12as cells used here were produced by Genome Engineering Oxford (GEO) as detailed in Materials and Methods. Purified nuclei were lysed and used for western blot analysis with antibodies against the N-terminus of Pol II, Pol II Ser2P, Pol II Ser5P, SPT5, pS666 and pT806 (Figure 5). The CDK9 inhibitor DRB was used as a control for loss of SPT5 phosphorylation caused by CDK9 inhibition and Lamin B was used as a loading control. This analysis determines the phosphorylation state of Pol II and SPT5 both on chromatin and in the nucleoplasm.

**Figure 5.**
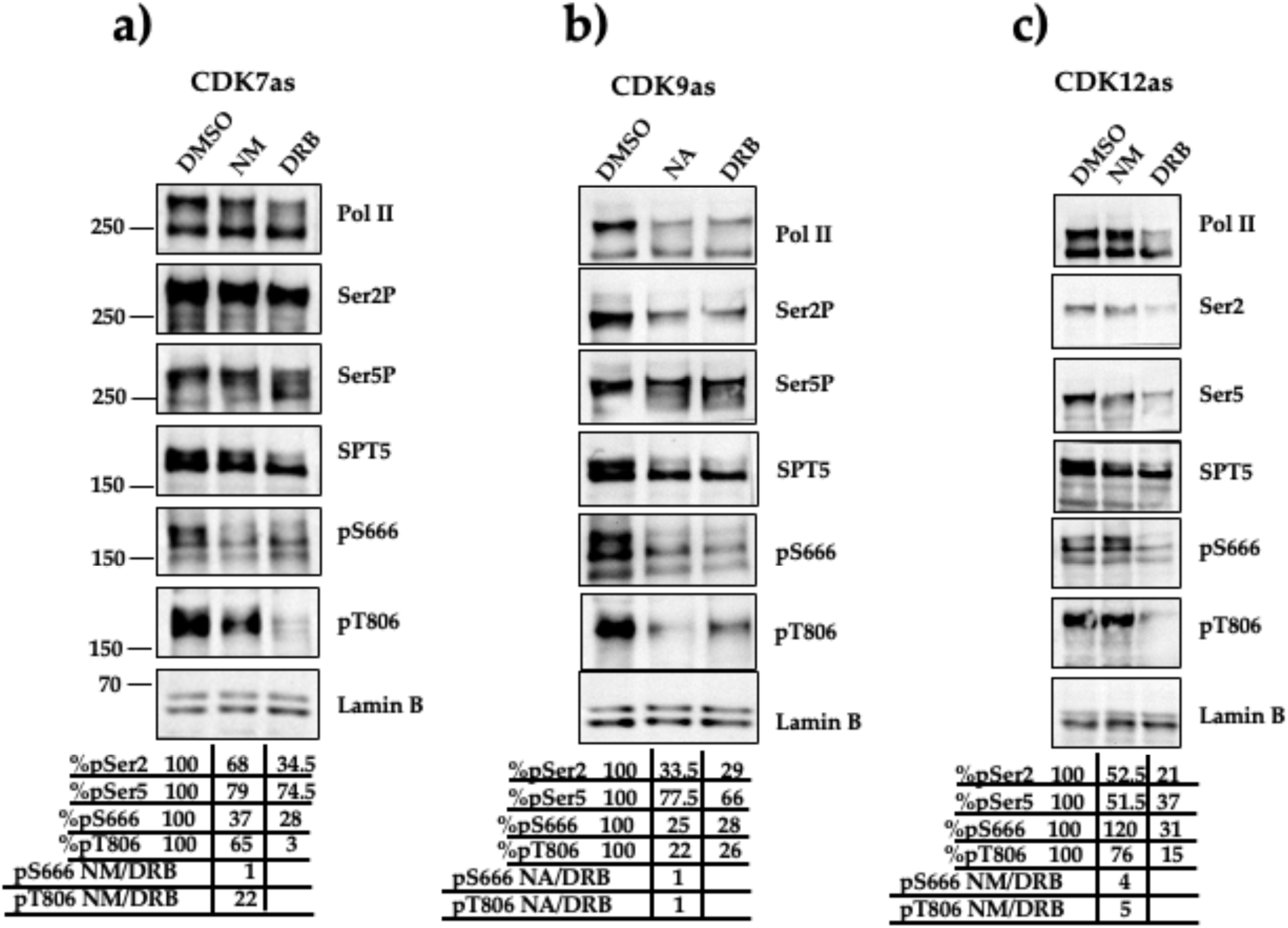
Inhibition of CDK7, CDK9 or CDK12 affects phosphorylation of SPT5 at S666 and T806. Western blot analysis of eluates from **a)** CDK7as cells treated with DMSO, 1-NM-PP1 or DRB as indicated at the top, **b)** CDK9as cells treated with DMSO, NA or DRB as indicated at the top, **c)** CDK12as cells treated with DMSO, NM or DRB as indicated at the top. Lamin B was used as an internal loading control and the averages from two biological repeats is noted at the bottom. DRB treatment of cells is used as a control for CDK9 inhibition in all cell lines.

Inhibition of CDK7as causes some reduction of Pol II phosphorylation at Ser5P, as expected, and a slight loss of Ser2P. It also reduces phosphorylation of S666 to 37% and of T806 to 65% relative to untreated samples (Figure 5a), suggesting a role for CDK7 in SPT5 phosphorylation, particularly of S666. This is in line with previous findings that inhibition of CDK7 using a specific small molecule inhibitor affects phosphorylation of S666[34]. The effect of CDK7as inhibition on pS666 is similar to the effect of DRB treatment, whereas DRB affects pT806 much more than inhibition of CDK7as, emphasizing the differential effect of inhibiting CDK7 on phosphorylation of these two residues.

Inhibition of CDK9as causes a loss of phosphorylation of Pol II at Ser2 and Ser5, as reported previously [38] (Figure 5b). In addition, pS666 and pT806 are reduced to 25% and 22%, respectively by CDK9as inhibition (Figure 5b). DRB has a very similar effect on phosphorylation of these residues. These results are in line with our previous findings that DRB causes loss of phosphorylation of SPT5[38].

In agreement with previous results, inhibition of CDK12as causes some reduction of both Pol II Ser2P and Ser5P[39]. CDK12 inhibition also causes a modest reduction of phosphorylation of T806 to 76%, whereas no loss of phosphorylation of S666 is detected, with rather an increase to 120% (Figure 5c). Treatment of these cells with DRB, instead, causes a much greater reduction of phosphorylation of both residues, as expected. Inhibition of CDK12as therefore has a modest negative effect only on T806 phosphorylation. However, phosphorylation of other residues may be affected and there appears to be a marked loss of the upper SPT5 band, which is likely to be the fully phosphorylated form.

These results indicate that inhibition of each of these three kinases can affect SPT5 phosphorylation. However, the biggest effect on phosphorylation of S666 and T806 is caused by inhibition of CDK9.

### Inhibition of CDK9 causes loss of association of PAF complex subunits with SPT5

PAF1C interacts with Pol II and SPT5 after phosphorylation of the Pol II CTD, NELF and SPT5 by CDK9[18]. Here, we have used ectopic expression of C-terminally FLAG-tagged full-length SPT5 (FLAG-SPT5) (Figure 1) to facilitate pulldown of SPT5 and western analysis of the associated proteins. We have used CDK9as cells transiently transfected with a plasmid encoding FLAG-SPT5 before and after treatment of the cells for 30 minutes with 1-NA-PP1 to determine the effect of loss of phosphorylation on the association of PAF1C subunits with SPT5. Nuclei were purified and lysed before incubating with anti-FLAG antibodies bound to IgG-bearing magnetic beads. After washing, the FLAG-tagged SPT5 and associated proteins were eluted by heating with LDS gel loading buffer with DTT, followed by western analysis (Figure 6). A plasmid encoding only the FLAG tag (Figure 1) was transfected as a control.

**Figure 6.**
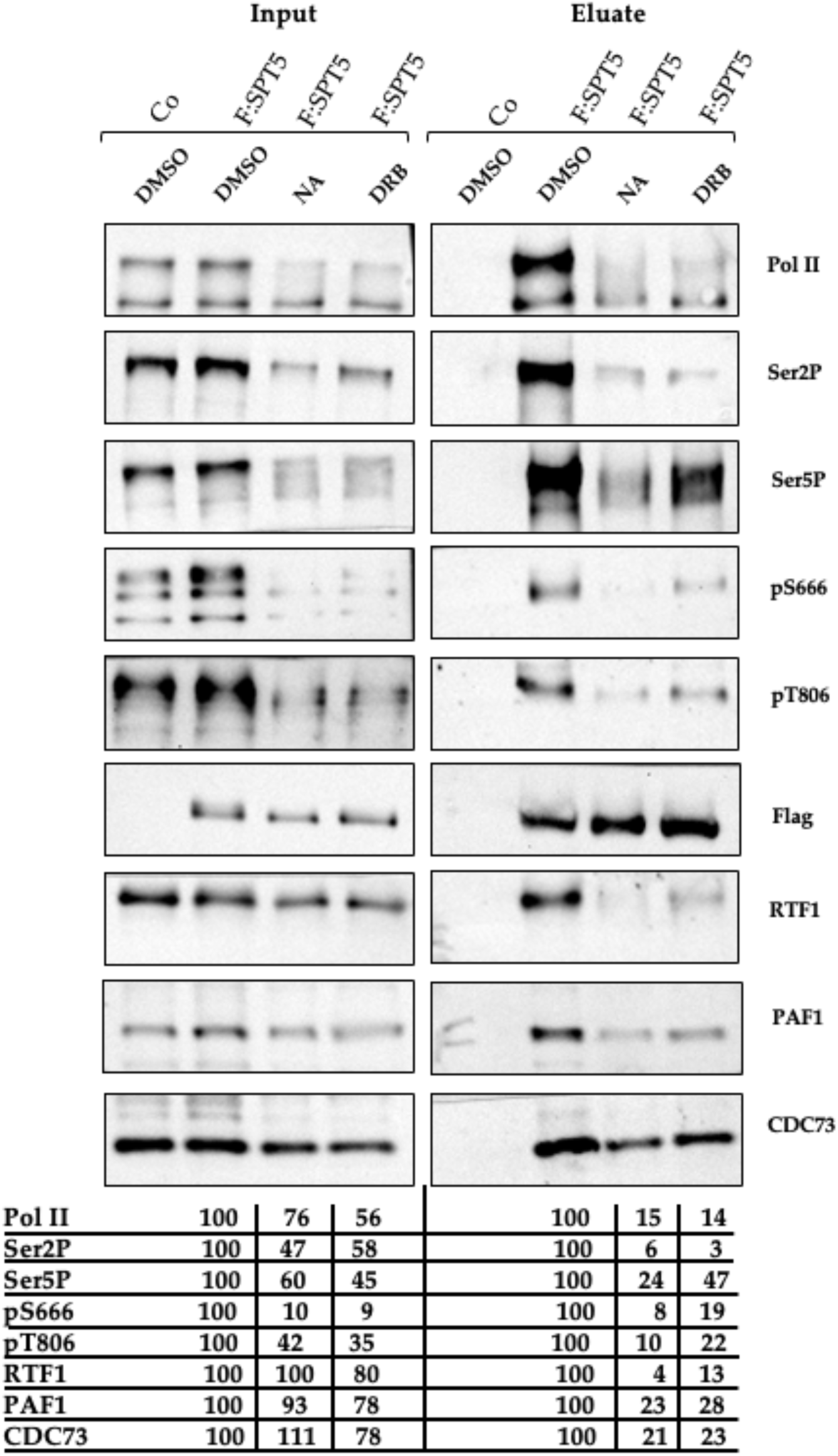
Inhibition of CDK9 causes loss of association of PAF complex subunits with SPT5. Western blot analysis of inputs and eluates of pulldown of C-terminally FLAG-tagged SPT5 (F:SPT5=FLAG-SPT5). The constructs transfected and the drug treatments are indicated at the top. CO=control template expressing only FLAG. NA= 1-NA-PP1. The antibodies used are indicated on the right and the levels are normalized to the level of FLAG detected with the anti-FLAG antibody and noted at the bottom as a percentage of the value for the cells transfected with FLAG and treated with DMSO.

Pol II and PAF complex subunits PAF1, RTF1 and CDC73 are pulled down with FLAG-SPT5, as expected according to previous results[12, 50]. Inhibition of CDK9as causes loss of phosphorylation of pS666 and pT806 on the pulled-down SPT5 to 8% and 10% of the control respectively, in line with the results in Figure 5b. The reduction of pT806 after CDK9 inhibition by 1-NA-PP1 in the input samples appears to be less marked (reduced to 42% of the control), possibly because the multiple bands reflecting both the endogenous and ectopically-expressed SPT5 complicate the quantitation. CDK9as inhibition also causes loss of association of the FLAG-SPT5 with PAF1 (reduced to 23% of the control versus 93% in the input), CDC73 (reduced to 21% of the control versus 111% in the input) and a marked effect on association of RTF1, which reduced to 4% of the control versus 100% in the input. There is also some loss of Pol II association (reduced to 15% of the control versus 76% in the input) and reduction of the ratio of Ser2P and Ser5P to Pol II relative to untreated samples and inputs. The results of DRB inhibition are similar.

These results confirm the association of SPT5 with Pol II and subunits of the PAF complex *in vivo* and indicate that specific inhibition of CDK9as causes loss of association of SPT5 with Pol II and PAF1C, with a particularly marked effect on the association of RTF1.

### Inhibition of CDK9 affects chromatin association of phosphorylated SPT5

To determine the effect of inhibition of CDK9as on the chromatin association of total SPT5 and SPT5 phosphorylated on either S666 or T806, we have carried out ChIP at the beginning of the model *KPNB1* gene (Figure 7a) using antibodies to total SPT5, pS666 and pT806 before and after inhibition of CDK9as for 30 minutes (Figure 7b).

**Figure 7.**
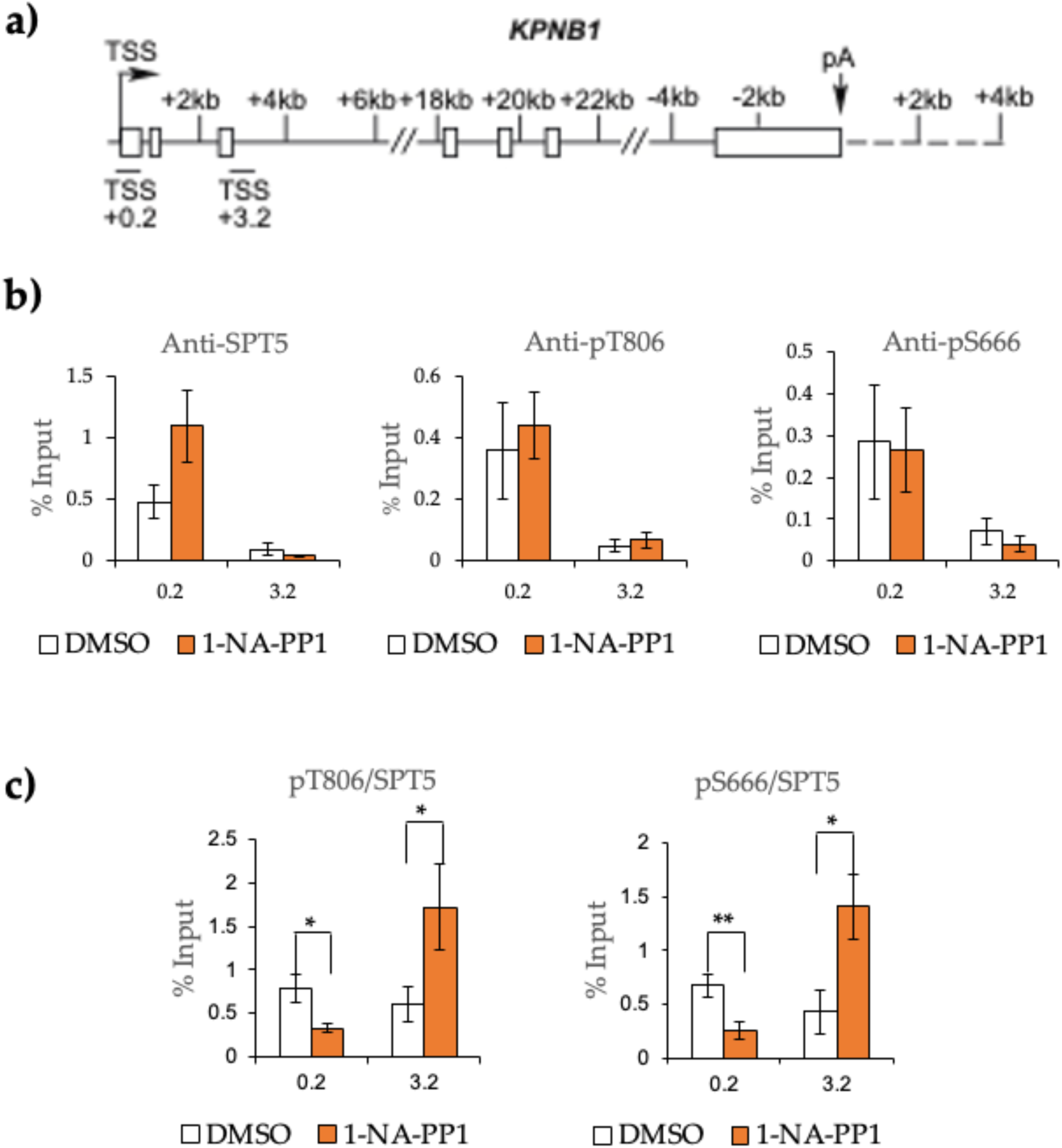
Inhibition of CDK9 affects chromatin association of phosphorylated SPT5. **a)** Schematic of KPNB1 with the position of primer sets used for chromatin immunoprecipitation (ChIP) shown. **b)** ChIP using CDK9as cells treated with DMSO or 1-NA-PP1 as indicated underneath each panel, using the antibodies indicated above. **c)** the ratio of pS666P or pT806 to SPT5. Error bars indicate standard error of the mean of 3 biological replicates. Asterisks show the statistical significance (Student’s unpaired t-test); *P < 0.05. **P<0.01.

In agreement with previous findings[29], SPT5, pS666 and pT806 are higher at the transcription start site (TSS) than at +3.2 kB of the KPNB1 gene before inhibition (Figure 7b, c). Inhibition of CDK9AS for 30 minutes causes an increase of the level of SPT5 at the beginning of KPNB1 and a reduction at +3.2 kB, which is where Pol II is reduced by inhibiting CDK9 for 30 minutes[38]. pS666 and pT806 relative to SPT5 are significantly decreased at the promoter region of KPNB1 after CDK9as inhibition (Figure 7c). However, both pS666 and pT806 relative to SPT5 significantly increase at +3.2 kB after CDK9as inhibition.

These results confirm that inhibiting CDK9 affects phosphorylation of SPT5 on chromatin in the linker region and CTR1. The reduction of pS666 and pT806 at the beginning of *KPNB1* by CDK9 inhibition is consistent with a role for phosphorylation of these regions early in the transcription cycle[29]. The increase in relative SPT5 phosphorylation at +3.2 kB may indicate that phosphorylated SPT5 is selectively retained when SPT5 and Pol II occupancy at this region is reduced by CDK9 inhibition.

## Discussion

SPT5 is a central player in elongation of transcription from yeast to man and its function is regulated by phosphorylation and dephosphorylation during the transcription cycle. Phosphorylation promotes interaction with the PAF1C elongation complex. Three distinct regions towards the C-terminus of SPT5, the linker between the KOW5 and KOW5 domains, CTR1 and CTR2, are reversibly phosphorylated *in vivo* and previous results implicate CDK7 and CDK9 in SPT5 phosphorylation[12, 34, 38, 46]. Here we have investigated the effect of *in vitro* phosphorylation of the C-terminus of SPT5 on interaction with members of PAF1C. The results confirm that the RTF1 subunit of PAF1C binds to the *in vitro* phosphorylated CTR1 alone and that this region of the SPT5 C-terminus is necessary for efficient interaction. We did not detect binding of other members of PAF1C to the *in vitro* phosphorylated C-terminus of SPT5. Although this may be due to technical problems, we cannot rule out that areas additional to the C-terminus of SPT5 or other proteins, for example Pol II, are necessary for interaction with other members of PAF1C *in vitro*. In addition, although RTF1 is generally considered to be a subunit of PAF1C, in metazoans, it is less stably associated with the complex than others and there are some indications that human RTF1 can function as an elongation factor without the rest of PAF1C[58, 59]. Thus, the requirements for interaction of SPT5 with the different PAF1C subunits may differ. Relevant to this, mutation of all potential phosphorylation sites in the linker, CTR1 and CTR2 regions affected chromatin association of RTF1 but not the PAF1 subunit of PAF1C[12], indicating that there are alternative pathways for PAF1 recruitment.

In *S. cerevisiae*, SPT5 is a smaller protein with a C-terminal region (CTR) comprising approximately 15 hexapeptide repeats with a consensus sequence of S(T/A)WGG(Q/A) rather than the GS(Q/R)TPXY consensus of the CTR1 repeats in human SPT5[60]. This CTR is phosphorylated by Bur1, a CDK9 homologue, and phosphorylation mediates the recruitment of the Paf1 complex (Paf1c), which is equivalent to human PAF1C[61], histone modifying enzymes and 3’ RNA-processing factors to elongating RNA polymerase II[60, 62–66]. As in humans, the yeast RTF1 homologue, Rtf1, plays a key role in the interaction between S. cerevisiae SPT5 and the Paf1 complex [65, 67] and both Paf1 and Rtf1 are lost from genes when the Spt5 CTR is deleted or mutated to eliminate phosphorylation sites[66]. In addition, depletion of PP1, which results in increased Spt5 phosphorylation, causes a termination defect due to Paf1c retention at the end of genes[66]. Thus, although the yeast SPT5 is less complex, with different CTR repeats, the control of elongation factor recruitment by SPT5 phosphorylation is an evolutionarily-conserved mechanism.

We have also used antibodies to phosphorylated residues within the linker between the KOW5 and KOW5 domains (pS666) and CTR1 (pT806) and HEK293 cell lines where the endogenous CDK7, CDK9 or CDK12 is analog-sensitive (as) to investigate the role of these CDKs in phosphorylating SPT5. Some loss of SPT5 phosphorylation occurs after individual inhibition of each of the three CDKs, although inhibiting CDK12as only causes a modest reduction of pT806 and rather increases pS666. CDK7as inhibition causes a greater reduction of pS666 than pT806, consistent with previous results identifying S666 and S671 as CDK7 targets after CDK7 inhibition with the covalent inhibitor SY-351[34]. CDK9as inhibition causes a marked reduction of both pS666 and pT806, indicating that it is a major SPT5 kinase, in agreement with previous work[2, 32, 37]. Notably, the effect of inhibiting each kinase is different, emphasizing their distinct roles in transcription. In addition, neither CDK7 nor CDK12 can compensate for loss of CDK9as activity, consistent with the role of CDK9 as a master transcription regulator[2, 32, 37].

CDK9as inhibition in cells has a marked effect on the interaction between SPT5 and PAF1C members; PAF1, CDC73 and RTF1 are all pulled down with FLAG-tagged full-length SPT5 and these interactions are markedly reduced when CDK9as is inhibited. These results confirm the major role of CDK9 in the recruitment of elongation factors to the elongating Pol II complex through SPT5 phosphorylation. PAF1C is involved in several transcriptional processes including promoting elongation of transcription through chromatin[68], regulating release of paused Pol II and helping to recruit CDK12[59, 69, 70]. SPT5 phosphorylation is therefore a critical step for efficient and productive elongation.

It is important to keep in mind that some of the effects on SPT5 phosphorylation of inhibiting CDK7as, CDK12as and CDK9as may be secondary to changes in the speed or processivity of Pol II and/or the recruitment/stability of the other kinases, making the order of events difficult to unpick. For example, CDK9as inhibition may affect SPT5 T806 phosphorylation through loss of CDK12 activity caused by loss of PAF1C association[70]. CDK12 inhibition causes loss of the PAF1C subunits Leo1 and CDC73 from transcribed genes [39] and this may be due to loss of phosphorylation of SPT5, Pol II [39] or Leo1[40]. Interestingly, CDC73 has been shown to activate the kinase activity of CDK12[71], highlighting the complex interactions between all of these elongation factors. In addition, although our analyses indicate that CDK12 inhibition has only a limited effect on T806 phosphorylation, phosphorylation of other SPT5 residues important for the association of PAF1C subunits may be affected by inhibiting CDK12. The increase of pS666 after CDK12 inhibition may be because Pol II is slower[39], giving CDK7 more time to phosphorylate SPT5. Furthermore, CDK9 inhibition affects initiation[72], which could affect CDK7 activity. It should also be kept in mind that these kinases function together with and regulate antagonistic phosphatases[48], which will contribute to the final levels of SPT5 phosphorylation at the two sites recognized by the antibodies in addition to other residues.

Our analyses therefore help to characterize the effect of mammalian SPT5 phosphorylation on association of PAF1C subunits *in vitro* and *in vivo* and the roles that CDK7, CDK9 and CDK12 play in SPT5 phosphorylation.

## Author Contributions

Conceptualization, S.M., I.M.; methodology, S.M., S.B., I.M.; investigation, S.M., S.S., S.B. I.M.; resources, S.M., S.B., J.R., G.R.M and I.M.; data curation, S.M and I.M.; writing—original draft preparation, S.M.; writing—review and editing, S.M. and I.M.; supervision, S.M.; project administration, S.M.; funding acquisition, S.M. All authors have seen and approved the manuscript, and it has not been accepted or published elsewhere.

## Distribution/reuse option

CC-BY-NC-ND

## Acknowledgements

The authors are very grateful to Professor Karen Arndt of the University of Pittsburgh for insightful comments on the manuscript and picking up entirely avoidable errors. Any errors that remain are the authors’ own.

## References

1. Yamaguchi, Y., H. Shibata, and H. Handa, Transcription elongation factors DSIF and NELF: promoter-proximal pausing and beyond. Biochim Biophys Acta, 2013. 1829(1): p. 98–104.

2. Fujinaga, K., F. Huang, and B.M. Peterlin, P-TEFb: The master regulator of transcription elongation. Mol Cell, 2023. 83(3): p. 393–403.

3. Buratowski, S. and K.M. Arndt, Controlling Transcription Elongation and Termination: More Than a Means to An End. J Mol Biol, 2025. 437(1): p. 168886.

4. Sun, R. and R.P. Fisher, The CDK9-SPT5 Axis in Control of Transcription Elongation by RNAPII. J Mol Biol, 2025. 437(1): p. 168746.

5. Hartzog, G.A. and J. Fu, The Spt4-Spt5 complex: a multi-faceted regulator of transcription elongation. Biochim Biophys Acta, 2013. 1829(1): p. 105–15.

6. Zaborowska, J., S. Egloff, and S. Murphy, The pol II CTD: new twists in the tail. Nat Struct Mol Biol, 2016. 23(9): p. 771–7.

7. Song, A. and F.X. Chen, The pleiotropic roles of SPT5 in transcription. Transcription, 2022. 13(1-3): p. 53–69.

8. Mooney, R.A., et al., NusG-Spt5 Transcription Factors: Universal, Dynamic Modulators of Gene Expression. J Mol Biol, 2025. 437(1): p. 168814.

9. Francette, A.M., et al., Evolutionary analysis of transcription elongation factors reveals conserved and lineage-specific regulatory domains. PLoS Biol, 2026. 24(6): p. e3003855.

10. Wada, T., et al., DSIF, a novel transcription elongation factor that regulates RNA polymerase II processivity, is composed of human Spt4 and Spt5 homologs. Genes Dev, 1998. 12(3): p. 343–56.

11. Ivanov, D., et al., Domains in the SPT5 protein that modulate its transcriptional regulatory properties. Mol Cell Biol, 2000. 20(9): p. 2970–83.

12. Sun, R. and R.P. Fisher, Tripartite phosphorylation of SPT5 by CDK9 times pause release and tunes elongation rate of RNA polymerase II. Mol Cell, 2025. 85(9): p. 1743–1759 e5.

13. Martinez-Rucobo, F.W., et al., Architecture of the RNA polymerase-Spt4/5 complex and basis of universal transcription processivity. EMBO J, 2011. 30(7): p. 1302–10.

14. Li, W., C. Giles, and S. Li, Insights into how Spt5 functions in transcription elongation and repressing transcription coupled DNA repair. Nucleic Acids Res, 2014. 42(11): p. 7069–83.

15. Meyer, P.A., et al., Structures and Functions of the Multiple KOW Domains of Transcription Elongation Factor Spt5. Mol Cell Biol, 2015. 35(19): p. 3354–69.

16. Qiu, Y. and D.S. Gilmour, Identification of Regions in the Spt5 Subunit of DRB Sensitivity-inducing Factor (DSIF) That Are Involved in Promoter-proximal Pausing. J Biol Chem, 2017. 292(13): p. 5555–5570.

17. Dollinger, R., et al., Assessment of the roles of Spt5-nucleic acid contacts in promoter proximal pausing of RNA polymerase II. J Biol Chem, 2023. 299(9): p. 105106.

18. Vos, S.M., et al., Structure of activated transcription complex Pol II-DSIF-PAF-SPT6. Nature, 2018. 560(7720): p. 607–612.

19. Core, L. and K. Adelman, Promoter-proximal pausing of RNA polymerase II: a nexus of gene regulation. Genes Dev, 2019. 33(15-16): p. 960–982.

20. Aoi, Y. and A. Shilatifard, Transcriptional elongation control in developmental gene expression, aging, and disease. Mol Cell, 2023. 83(22): p. 3972–3999.

21. Moorefield, B., SPT5 roles in transcriptional elongation. Nat Struct Mol Biol, 2021. 28(10): p. 778.

22. Aoi, Y., et al., SPT5 stabilization of promoter-proximal RNA polymerase II. Mol Cell, 2021. 81(21): p. 4413–4424 e5.

23. Hu, S., et al., SPT5 stabilizes RNA polymerase II, orchestrates transcription cycles, and maintains the enhancer landscape. Mol Cell, 2021. 81(21): p. 4425–4439 e6.

24. Lindstrom, D.L., et al., Dual roles for Spt5 in pre-mRNA processing and transcription elongation revealed by identification of Spt5-associated proteins. Mol Cell Biol, 2003. 23(4): p. 1368–78.

25. Baejen, C., et al., Genome-wide Analysis of RNA Polymerase II Termination at Protein-Coding Genes. Mol Cell, 2017. 66(1): p. 38–49 e6.

26. Parua, P.K., et al., A Cdk9-PP1 switch regulates the elongation-termination transition of RNA polymerase II. Nature, 2018. 558(7710): p. 460–464.

27. Cortazar, M.A., et al., Control of RNA Pol II Speed by PNUTS-PP1 and Spt5 Dephosphorylation Facilitates Termination by a “Sitting Duck Torpedo” Mechanism. Mol Cell, 2019. 76(6): p. 896–908 e4.

28. Eaton, J.D., et al., A unified allosteric/torpedo mechanism for transcriptional termination on human protein-coding genes. Genes Dev, 2020. 34(1-2): p. 132–145.

29. Parua, P.K., et al., Distinct Cdk9-phosphatase switches act at the beginning and end of elongation by RNA polymerase II. Nat Commun, 2020. 11(1): p. 4338.

30. Malumbres, M., Cyclin-dependent kinases. Genome Biol, 2014. 15(6): p. 122.

31. Parua, P.K. and R.P. Fisher, Dissecting the Pol II transcription cycle and derailing cancer with CDK inhibitors. Nat Chem Biol, 2020. 16(7): p. 716–724.

32. Pluta, A.J., et al., Cyclin-dependent kinases: Masters of the eukaryotic universe. Wiley Interdiscip Rev RNA, 2023. 15(1): p. e1816.

33. Fisher, R.P., Cdk7: a kinase at the core of transcription and in the crosshairs of cancer drug discovery. Transcription, 2019. 10(2): p. 47–56.

34. Rimel, J.K., et al., Selective inhibition of CDK7 reveals high-confidence targets and new models for TFIIH function in transcription. Genes Dev, 2020. 34(21-22): p. 1452–1473.

35. Velychko, T., et al., CDK7 kinase activity promotes RNA polymerase II promoter escape by facilitating initiation factor release. Mol Cell, 2024. 84(12): p. 2287–2303 e10.

36. Peterlin, B.M. and D.H. Price, Controlling the elongation phase of transcription with P-TEFb. Mol Cell, 2006. 23(3): p. 297–305.

37. Bres, V., S.M. Yoh, and K.A. Jones, The multi-tasking P-TEFb complex. Curr Opin Cell Biol, 2008. 20(3): p. 334–40.

38. Tellier, M., et al., CDK9 and PP2A regulate RNA polymerase II transcription termination and coupled RNA maturation. EMBO Rep, 2022. 23(10): p. e54520.

39. Tellier, M., et al., CDK12 globally stimulates RNA polymerase II transcription elongation and carboxyl-terminal domain phosphorylation. Nucleic Acids Res, 2020. 48(14): p. 7712–7727.

40. Qiu, M., et al., CDK12 and Integrator-PP2A complex modulates LEO1 phosphorylation for processive transcription elongation. Sci Adv, 2023. 9(20): p. eadf8698.

41. Blazek, D., et al., The Cyclin K/Cdk12 complex maintains genomic stability via regulation of expression of DNA damage response genes. Genes Dev, 2011. 25(20): p. 2158–72.

42. Dubbury, S.J., P.L. Boutz, and P.A. Sharp, CDK12 regulates DNA repair genes by suppressing intronic polyadenylation. Nature, 2018. 564(7734): p. 141–145.

43. Fan, Z., et al., CDK13 cooperates with CDK12 to control global RNA polymerase II processivity. Sci Adv, 2020. 6(18).

44. Yamada, T., et al., P-TEFb-mediated phosphorylation of hSpt5 C-terminal repeats is critical for processive transcription elongation. Mol Cell, 2006. 21(2): p. 227–37.

45. Laitem, C., et al., CDK9 inhibitors define elongation checkpoints at both ends of RNA polymerase II-transcribed genes. Nat Struct Mol Biol, 2015. 22(5): p. 396–403.

46. Sanso, M., et al., P-TEFb regulation of transcription termination factor Xrn2 revealed by a chemical genetic screen for Cdk9 substrates. Genes Dev, 2016. 30(1): p. 117–31.

47. Vervoort, S.J., et al., The PP2A-Integrator-CDK9 axis fine-tunes transcription and can be targeted therapeutically in cancer. Cell, 2021. 184(12): p. 3143–3162 e32.

48. Cossa, G., et al., Protein phosphatases in the RNAPII transcription cycle: erasers, sculptors, gatekeepers, and potential drug targets. Genes Dev, 2021. 35(9-10): p. 658–676.

49. Fong, N., et al., The pausing zone and control of RNA polymerase II elongation by Spt5: Implications for the pause-release model. Mol Cell, 2022. 82(19): p. 3632–3645 e4.

50. Wier, A.D., et al., Structural basis for Spt5-mediated recruitment of the Paf1 complex to chromatin. Proc Natl Acad Sci U S A, 2013. 110(43): p. 17290–5.

51. Schafer, F., et al., Purification of GST-Tagged Proteins. Methods Enzymol, 2015. 559: p. 127–39.

52. Dignam, J.D., R.M. Lebovitz, and R.G. Roeder, Accurate transcription initiation by RNA polymerase II in a soluble extract from isolated mammalian nuclei. Nucleic Acids Res, 1983. 11(5): p. 1475–89.

53. Nojima, T., et al., Mammalian NET-seq analysis defines nascent RNA profiles and associated RNA processing genomewide. Nat Protoc, 2016. 11(3): p. 413–28.

54. Laitem, C., et al., CTCF regulates NELF, DSIF and P-TEFb recruitment during transcription. Transcription, 2015. 6(5): p. 79–90.

55. Renaud, J.B., et al., Improved Genome Editing Efficiency and Flexibility Using Modified Oligonucleotides with TALEN and CRISPR-Cas9 Nucleases. Cell Rep, 2016. 14(9): p. 2263–2272.

56. Abramson, J., et al., Accurate structure prediction of biomolecular interactions with AlphaFold 3. Nature, 2024. 630(8016): p. 493-500.

57. Goddard, T.D., et al., UCSF ChimeraX: Meeting modern challenges in visualization and analysis. Protein Sci, 2018. 27(1): p. 14–25.

58. Cao, Q.F., et al., Characterization of the Human Transcription Elongation Factor Rtf1: Evidence for Nonoverlapping Functions of Rtf1 and the Paf1 Complex. Mol Cell Biol, 2015. 35(20): p. 3459–70.

59. Francette, A.M., S.A. Tripplehorn, and K.M. Arndt, The Paf1 Complex: A Keystone of Nuclear Regulation Operating at the Interface of Transcription and Chromatin. J Mol Biol, 2021. 433(14): p. 166979.

60. Zhou, K., et al., Control of transcriptional elongation and cotranscriptional histone modification by the yeast BUR kinase substrate Spt5. Proc Natl Acad Sci U S A, 2009. 106(17): p. 6956–61.

61. Betz, J.L., et al., Phenotypic analysis of Paf1/RNA polymerase II complex mutations reveals connections to cell cycle regulation, protein synthesis, and lipid and nucleic acid metabolism. Mol Genet Genomics, 2002. 268(2): p. 272–85.

62. Liu, Y., et al., Phosphorylation of the transcription elongation factor Spt5 by yeast Bur1 kinase stimulates recruitment of the PAF complex. Mol Cell Biol, 2009. 29(17): p. 4852–63.

63. Ding, B., D. LeJeune, and S. Li, The C-terminal repeat domain of Spt5 plays an important role in suppression of Rad26-independent transcription coupled repair. J Biol Chem, 2010. 285(8): p. 5317–26.

64. Mayer, A., et al., The spt5 C-terminal region recruits yeast 3’ RNA cleavage factor I. Mol Cell Biol, 2012. 32(7): p. 1321–31.

65. Mayekar, M.K., R.G. Gardner, and K.M. Arndt, The recruitment of the Saccharomyces cerevisiae Paf1 complex to active genes requires a domain of Rtf1 that directly interacts with the Spt4-Spt5 complex. Mol Cell Biol, 2013. 33(16): p. 3259–73.

66. Namjilsuren, S. and K.M. Arndt, Glc7/PP1 triggers Paf1 complex dissociation from RNA polymerase II to enable transcription termination. Genes Dev, 2026. 40(9-10): p. 699–716.

67. Squazzo, S.L., et al., The Paf1 complex physically and functionally associates with transcription elongation factors in vivo. EMBO J, 2002. 21(7): p. 1764–74.

68. Kim, J., M. Guermah, and R.G. Roeder, The human PAF1 complex acts in chromatin transcription elongation both independently and cooperatively with SII/TFIIS. Cell, 2010. 140(4): p. 491–503.

69. Chen, F.X., et al., PAF1, a Molecular Regulator of Promoter-Proximal Pausing by RNA Polymerase II. Cell, 2015. 162(5): p. 1003–15.

70. Yu, M., et al., RNA polymerase II-associated factor 1 regulates the release and phosphorylation of paused RNA polymerase II. Science, 2015. 350(6266): p. 1383–6.

71. Lopez Martinez, D., et al., PAF1C-mediated activation of CDK12/13 kinase activity is critical for CTD phosphorylation and transcript elongation. Mol Cell, 2025. 85(10): p. 1952–1967 e8.

72. Gressel, S., et al., CDK9-dependent RNA polymerase II pausing controls transcription initiation. Elife, 2017. 6.

