## Supplemental figure for "Inhibition of CDK7, CDK9 or CDK12 affects phosphorylation of human SPT5"

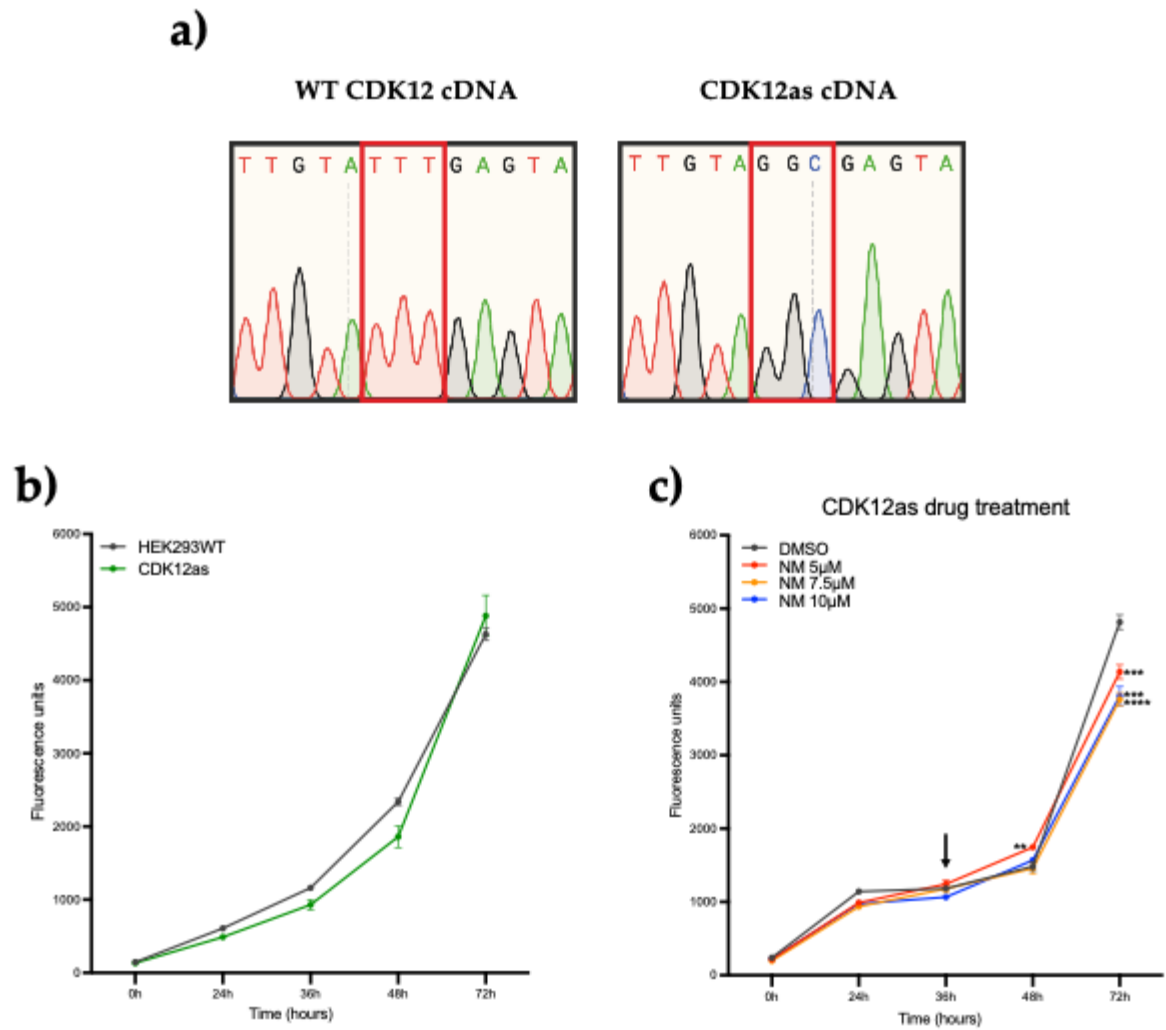

### Supplementary Figure. Characterization of CDK12as cells

**a)** Sanger sequencing traces of sequencing the cDNA of CDK12 from wild type (WT) 293 cells or CDK12as cells as detailed in [39]. **b)** Growth curve of HEK293 wild type (HEK293WT) and CDK12as measured as detailed in Materials and Methods. **c)** Growth curve of CDK12as with DMSO or a range of concentrations of 1-NM-PP1 added at 36 hours as indicated by the vertical arrow.
